# Novel extracellular vesicle release pathway facilitated by toxic superoxide dismutase 1 oligomers

**DOI:** 10.1101/2025.04.07.647611

**Authors:** Brianna Hnath, Nikolay V. Dokholyan

## Abstract

Amyotrophic lateral sclerosis (ALS) is a devastating neurodegenerative disease resulting in paralysis and death within three to five years. Mutations in over forty different proteins have been linked to ALS, leading to controversy whether ALS is one disease or many diseases with a similar phenotype. Mutations in Cu,Zn superoxide dismutase 1 (SOD1) are only found in 2-3% of ALS cases, yet misfolded SOD1 is found in both sporadic (sALS) and familial (fALS) patients. Yet, mutations in TDP-43 or FUS increase the level of misfolded SOD1 on extracellular vesicles (EVs). Additionally, small EVs isolated from ALS patient samples caused cell death of wild type motor neurons and myotubules. The toxicity and protein alterations of ALS EVs have led to the theory that EVs are responsible for the spread of ALS. We hypothesize that previously-identified toxic trimeric SOD1 is spreading on EVs in ALS and altering the spread of other ALS-related proteins, linking them to a common mechanism. To test our hypothesis, we isolate EVs from motor neuron-like cells expressing trimer stabilizing mutations and perform a sandwich enzyme-linked immunoassay (ELISA) (CD9 capture antibody) to quantify whether misfolded SOD1 and 17 other ALS-related proteins increase or decrease on EVs with trimer stabilization. We identify which EV release pathway is being affected by trimeric SOD1 utilizing endocytosis and exocytosis inhibitors, and determine if any specific EV-related proteins are altered with trimer stabilization. We establish that VAPB, VCP, and Stathmin-2 increase on EVs with trimer stabilization. The common pathway between SOD1 and three other ALS-associated proteins is affected by multiple pathways, including the Caveolae endocytosis pathway, suggesting a novel hybrid pathway of EV release present in ALS.

## Introduction

Over 5,000 people are diagnosed with ALS each year. Only 10% of these cases are familial, while the remaining 90% are sporadic cases^1,2^. Similar proteins are found to be misfolded in both fALS and sALS. Since 1993, mutations in more than forty different genes have been linked to ALS, and more continue to be identified^3^. The majority of the genetic mutations associated with ALS fall into four main functional categories: RNA processing, protein trafficking, cytoskeletal/axonal dynamics, and mitochondrial function^4^ (**Figure 1**). SOD1 is distinctive in that misfolded SOD1 has been shown to impact processes across all four categories. Many ALS-related proteins form large, insoluble aggregates when misfolded. Initially, large aggregates were thought to transmit extracellularly, spreading ALS between cells^5^. Many studies focus on the potential prion-like misfolded propagation of ALS-related proteins without proposing one common mechanism of spread^5,6^. Recently, smaller soluble forms of disease-associated proteins have been identified as toxic, while the larger insoluble aggregates are considered protective to cells^7^. As these soluble misfolded proteins are detected in or on EVs, we posit that EVs may play a crucial role in the spread of ALS and this theory is becoming more prevalent^8^.

**Figure 1.**
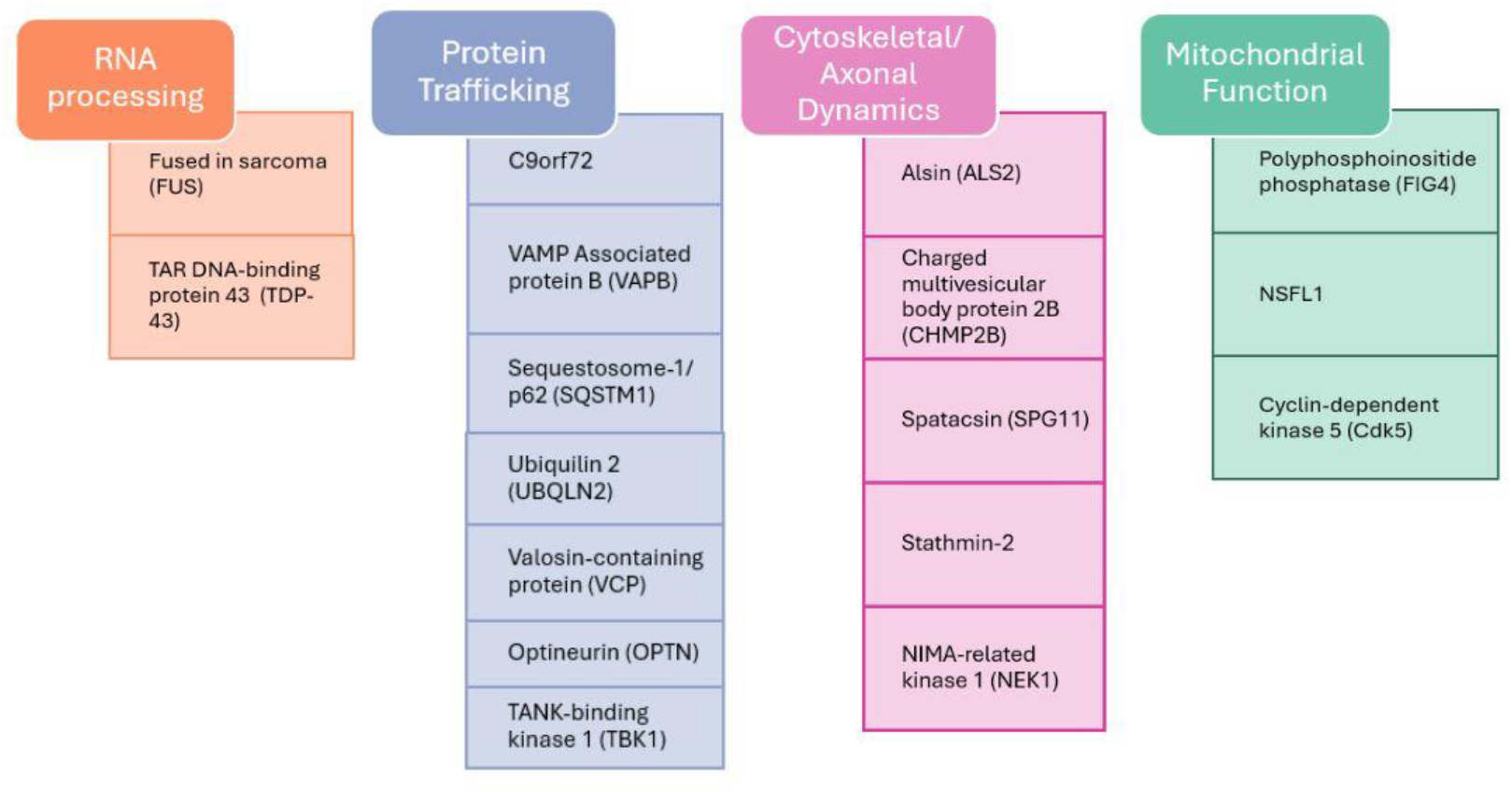
ALS-associated proteins can be divided into four groups based on their function.

SOD1 is a cytosolic antioxidant enzyme that naturally forms a dimer and converts superoxide radicals into oxygen and hydrogen peroxide^4^. Mutations in SOD1 destabilize the dimer interface, leading to the loss of metals and aggregation into both soluble and insoluble oligomers^9^. Mutant SOD1 (G93A) mouse models exhibit a loss of motor function, while SOD1 knockdown mouse models do not demonstrate the same phenotype^10,11^. This study led to the conclusion that SOD1 misfolding results in a toxic gain of function rather than a loss of function. While mutations in SOD1 are found in only 1-3% of all patients, misfolded SOD1 has been observed among patients with other ALS-related mutations (such as fused in sarcoma (FUS) and TDP-43) as well as in sporadic patients^12–15^. Several other factors beyond mutations cause SOD1 to destabilize and aggregate, including post-translational modifications (primarily glutathionylation)^16–18^, loss of metals^9,19^, crowding from overexpression^4^, and environmental toxins (such as BMAA and ammonia)^9,20–23^. The presence of misfolded SOD1 in sporadic and non-SOD1 familial ALS patients has been highly debated over the past decade, likely due to inconsistencies in the antibodies used^14,24,25^ and variations in tissue types and preparations. Forsberg, Pare, Grad, and Pokrishevsky all observed misfolded SOD1 in non-SOD1 fALS and sALS patient spinal cord sections using either immunohistochemistry or immunoprecipitation^14,15,26,27^. The difficulty of antibodies in identifying smaller oligomers of SOD1 has sustained the failed theory that large insoluble aggregates are responsible for the progression of ALS^23,28^.

When the SOD1 dimer interface is disrupted, SOD1 aggregates and forms a wide range of soluble oligomers and insoluble fibrils^9,18,19,29^. Most previous studies consider misfolded SOD1 to mean a mix of all aggregate sizes, yet the different-size oligomers have drastically independent toxicities and structures. In 2016, Proctor et al. determined that small soluble trimeric oligomers of SOD1 had the highest toxicity in cell models and that the toxicity correlated with the amount of thermodynamic stability of trimeric SOD1^30^. Zhu et al. further demonstrated that larger SOD1 oligomers and insoluble fibrils are protective to cells^28^. In 2022, Hnath and Dokholyan studied the aggregation mechanism of SOD1 using structural characterization assays and determined trimeric SOD1 is a structurally independent species that forms in direct competition with larger aggregates, as opposed to trimeric SOD1 being a preliminary step in larger aggregate formation^23^. Due to aggregate structural differences very few anti-misfolded SOD1 antibodies bind to trimeric SOD1, C4F6 is the only antibody we have identified that recognizes trimeric SOD1^24,29,30^. The inability of most antibodies to detect trimeric SOD1 may account for the drastic inconsistencies between studies quantifying misfolded SOD1 in patient samples.

The mechanism of spread of ALS throughout the body is largely unknown, EV release is increasingly recognized as a key mechanism in ALS pathogenesis, particularly in the spread of pathological proteins and disease-associated toxicity between cells^8,31^. EV biogenesis involves intricate endocytic and exocytic pathways that regulate cargo uptake, processing, and secretion, and dysfunction in these pathways may underlie the propagation of misfolded proteins such as SOD1 in ALS. Normally, intracellular proteins destined for EVs originate from the ER-Golgi network or are internalized through caveolin-dependent endocytosis, clathrin-dependent endocytosis, or pinocytosis. The proteins then traffic through early and late endosomes before being sorted for either lysosomal degradation or secretion^32^. However, in ALS, impaired endolysosomal processing can shift this balance, leading to excessive secretion of neurotoxic proteins on EVs rather than degradation^8^.

Misfolded SOD1 has been identified on the surface of extracellular vesicles (EVs) secreted from different ALS cell models (HEK293, Neuro2a, or NSC-34)^33–35^ or from ALS mouse models^34,36^. Misfolded SOD1 was found colocalized with neurosecretory vesicle proteins chromogranin A and B, indicating that misfolded SOD1 is secreted by cells instead of being released solely through apoptosis^34^. Small EVs from ALS patients were observed to be neurotoxic when added to stem cell-derived motor neuron and muscle cell cultures, yet larger EVs had no effect on viability^37^, suggesting the deadly progression of ALS may be vesicle size dependent. Overexpression of SOD1 mutants may impair ER-Golgi transport leading to Golgi fragmentation and ER stress^38,39^. Rab guanosine triphosphatases (GTPases) regulate all membrane trafficking events, and Rab5, Rab7, and Rab11 have been implicated in ALS^40^. Rab7 and Rab11 co-localize with the ALS-associated protein C9orf72^41^, which is expected since Rab7 controls endo-lysosomal transport and Rab11 regulates recycling endosomes—both essential for clearing misfolded proteins^42^. Mutations in ALS-associated Alsin (ALS2) disrupt Rab5 activation, impairing endosomal trafficking and vesicle biogenesis, which Rab5 normally regulates by facilitating early endosome fusion and cargo sorting^43^. Consequently, enlarged Rab5-positive endosomes are observed in spinal ALS motor neurons^44^. Understanding these dysregulated pathways could uncover new therapeutic targets aimed at disrupting the intercellular spread of ALS pathology. Beyond ALS2, multiple mutated genes found in ALS patients are involved in vesicular trafficking and have been shown to affect or coincide with SOD1 misfolding^45^. These genes can be grouped into four main functional categories as mentioned previously^4^: RNA processing—fused in sarcoma (FUS)^26^ and TAR DNA-binding protein 43 (TDP-43)^26^; protein trafficking –C9orf72^15^, vesicle-associated membrane protein-associated protein B (VAPB)^15,46^, Sequestosome-1/p62 (SQSTM1)^47,48^, Ubiquilin 2 (UBQLN2)^49^, Valosin-containing protein (VCP)^50^, Optineurin (OPTN), and TANK-binding kinase 1 (TBK1)^51^; cytoskeletal/axonal dynamics – ALS2^15,48,52^, charged multivesicular body protein 2B (CHMP2B), Spatacsin (SPG11), Stathmin-2^38^, and NIMA-related kinase 1 (NEK1); and mitochondrial function – polyphosphoinositide phosphatase (FIG4), NSFL1, and cyclin-dependent kinase 5 (Cdk5) (**Figure 1**). We propose that misfolded, trimeric, SOD1 is a central component of the mechanism of spreading of ALS on EVs, offering novel therapeutic targets to alter the disease progression.

To test our hypothesis and elucidate the pathways involved in the spread of toxic SOD1 and other ALS-associated proteins on EVs, we established a system to test the effect of SOD1 trimer on other proteins that do not solely rely on the use of antibodies (due to the difficulty for most antibodies to bind to trimeric SOD1). We used previously designed mutations that stabilize the trimeric form of SOD1 as well as the common disease mutation of A4V which partially stabilizes trimeric SOD1 and determined that stabilizing trimeric SOD1 leads to an increase in misfolded SOD1 on EVs. We used this method to test whether the stabilization of the SOD1 trimer affects the levels of a panel of other ALS-associated proteins on EVs. Upon narrowing the panel down to three main proteins (VAPB, VCP, and Stathmin-2), we determined the effect of six different exocytosis and endocytosis inhibitors on the levels of VAPB, VCP, and Stathmin-2 with trimer stabilization. Additionally, we tested the effect of trimer stabilization on a panel of twenty-two different EV-related proteins. We determined a novel hybrid pathway responsible for the release of altered EVs with trimeric SOD1 stabilization, shedding light on molecular events driving the disease propagation.

## Results

### Trimeric SOD1 stabilization increases misfolded SOD1 on the surface of Evs

We hypothesize that cytotoxic trimeric SOD1 is spreading on EVs throughout ALS and altering the spread of other ALS-related proteins. Grad et al. have previously determined that misfolded SOD1 is present on the exterior of vesicles secreted by cells stably expressing WT or mutant SOD1^27,33^, but the conformation of misfolded SOD1 and other proteins being altered on EVs when SOD1 is misfolded has not been established. We exploited mutations that progressively increase trimer stabilization (A4V mildly stabilizes trimer formation, and F20L/H46Q (FH) strongly stabilizes trimer formation)^23^ to determine if stabilizing trimeric SOD1 increases the concentration of misfolded SOD1 on EVs. The antibody C4F6 is the only antibody that binds trimeric SOD1^24,30^, but it also binds other misfolded SOD1 species, making it difficult to isolate molecular effects due solely to SOD1 trimers. Incrementally stabilizing trimeric SOD1 in cells circumvents this difficulty. We previously designed FH mutant SOD1 molecule that promotes the stabilization of SOD1 in a toxic trimeric state^23^ (**Figure 2A,B**). Overexpression of FH in NSC-34 cells leads to a significant increase of misfolded SOD1 (anti-C4F6 antibody) concentration on EVs, compared to overexpression of wild type (WT) or a commonly found SOD1 mutation (A4V) (**Figure 2C**). Due to the previously established tendency for the FH mutations to promote trimer formation^23^, it is likely that the misfolded species being identified by C4F6 are trimeric SOD1. We utilize this system to determine pathways responsible for the spread of toxic trimeric SOD1 EVs.

**Figure 2.**
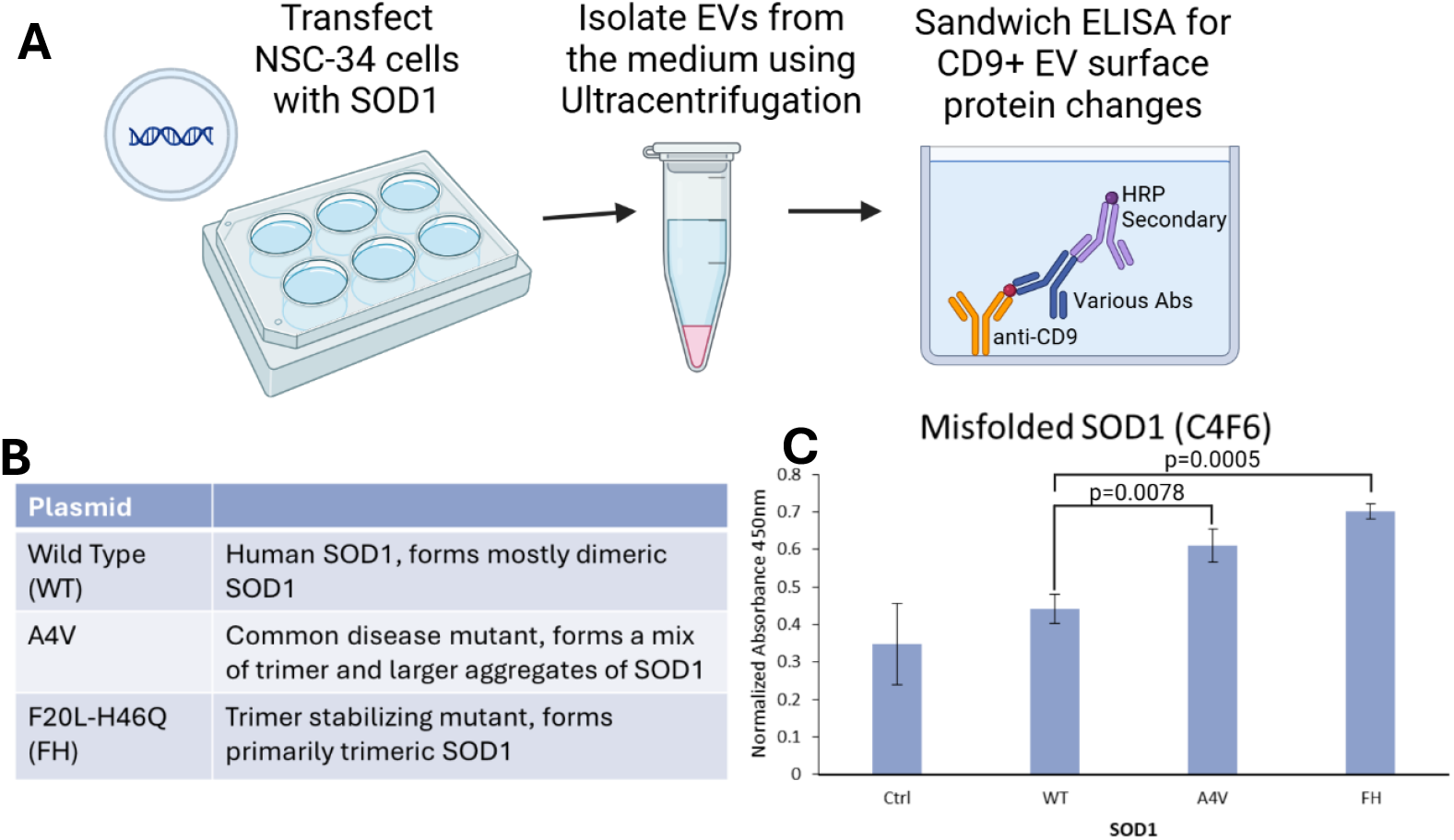
Stabilization of trimeric SOD1. A) Schematic diagram illustrating the workflow of transfecting NSC-34 cells with SOD1, isolating extracellular vesicles (EVs) from the culture medium via ultracentrifugation, and analyzing CD9+ EV surface protein changes using a sandwich ELISA. B) SOD1 plasmid mutations (or WT) to incrementally stabilize trimeric SOD1 in cells. C) Incrementally stabilizing trimeric SOD1 significantly increased the level of misfolded SOD1 on EVs isolated through an anti-CD9 sandwich ELISA. WT to A4V p-value= 0.0078, WT to FH p-value=0.0005.

### ALS-associated proteins increase and decrease on EVs with trimeric SOD1 stabilization

Many ALS-associated proteins are already known to be linked to EV formation and release^45^. We selected a panel of 17 ALS-associated proteins that either had known associations with EVs or were the most commonly found ALS mutations (C9ORF72, TDP-43, and FUS) and used our incremental trimer stabilization system to determine if any ALS-associated proteins are altered on EVs due to trimeric SOD1. Concentrations of VAPB (p-value 0.009), VCP (p-value 0.002), and Stathmin-2 (p-value 0.008) increased on EVs from cells expressing trimeric SOD1(FH) compared to WT controls, while concentrations of SPG11 (p-value 0.017) and OPTN (not significant) decreased with trimer stabilization (**Figure 3**). We did not observe significant changes in the concentration of FUS, TDP-43, C9orf72, SQSTM1, UBQLN2, TBK1, ALS2, CHMP2B, NEK1, FIG4, NSFL1, and Cdk5 with trimer stabilization. Since VAPB, VCP, and Stathmin-2 are all known to be involved in EVs^45,53^, we chose to focus on these three proteins for the remainder of the study while still taking into account all significant protein changes when considering potential EV release pathways. VAPB is a transmembrane protein involved in the tethering of the endoplasmic reticulum (ER) to other organelles, as well as the regulation of lipids^54,55^, VCP is broadly involved in vesicle trafficking and protein degradation^56^, and Stathmin-2 is the microtubule-associated protein responsible for destabilizing microtubules, allowing for axonal repair and maintenance, and has also been associated with Golgi fragmentation^38,53^. All three proteins are integral to different parts of EV formation and release, making it difficult to narrow down to a singular common pathway. We additionally tested whether ALS-associated mutations in VAPB and VCP caused the same increase in the proteins on EVs, but the mutants had no significant effects (**Supplementary Figure S3**) (Stathmin-2 does not yet have any direct mutations linked to ALS, Stathmin-2 can undergo cryptic splicing by other ALS-associated proteins leading to a loss of function^57^). Next, we determine a common pathway of release for these three altered proteins.

**Figure 3.**
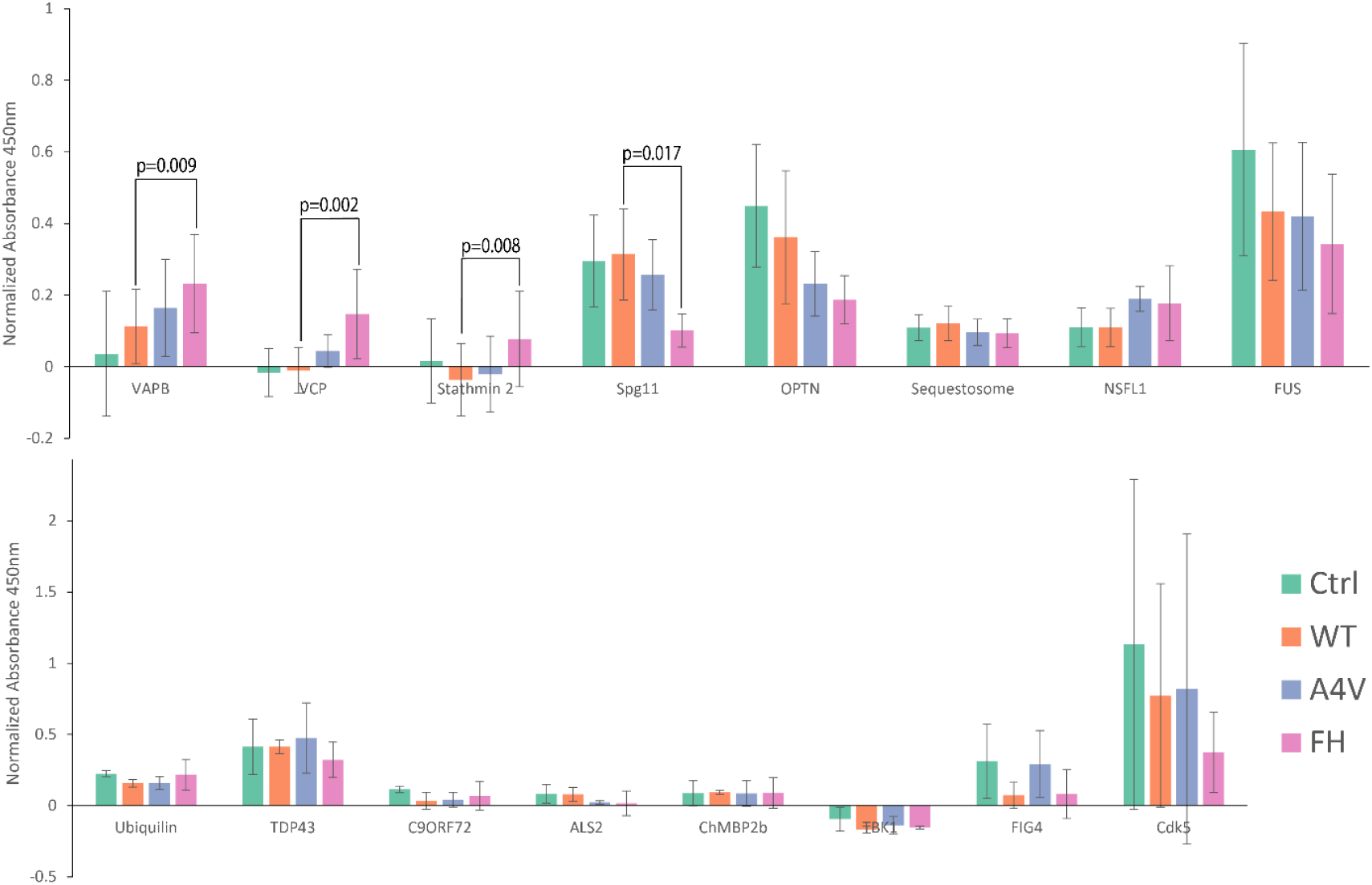
Stabilizing trimeric SOD1 alters ALS-associated protein levels on EVs. VAPB (p-value= 0.009), VCP (p-value=0.002), and Stathmin-2 (p-value=0.008) levels significantly increase on CD9+ EVs with trimeric SOD1 stabilization (FH). Spg11 (p-value= 0.017) levels significantly decrease, and OPTN levels appear to decrease although not significantly (p-value= 0.0546).

### Isolating the mechanism of release of altered Evs

To identify the pathway involved in the release of altered EVs due to trimer stabilization, we utilized two different methods. First, we used a panel of common endocytosis and exocytosis inhibitors^58,59^ to determine if inhibiting one pathway would affect the levels of VAPB, VCP, and Stathmin-2. Second, we evaluated groups of proteins involved in vesicle docking/release, ER/Golgi transport, movement of vesicles by the cytoskeleton, and lysosomal sorting to further narrow down the common pathway involved^60^.

Six different pathway inhibitors were tested; ESCRT-dependent exocytosis inhibitor Manumycin A, ESCRT-independent exocytosis inhibitor GW4869, macropinocytosis inhibitor Wortmannin, Na/H channel inhibitor (also a factor in macropinocytosis) ethyl isopropyl amiloride (EIPA), tyrosine kinase inhibitor (Caveolae-mediated endocytosis) Genistein, dynamin inhibitor MitMAB (affects both Caveolin-dependent and Clathrin-dependent endocytosis), and a clathrin endocytosis inhibitor Chlorpromazine were tested in NSC-34 cells (**Figure 4**)^58,59^. If a certain pathway was involved in the release of the trimer-altered EVs, we expected to observe a decrease in VAPB, VCP, and Stathmin-2 when the release pathway is inhibited. Instead, the only inhibitor that caused a significant change in FH for all three proteins was EIPA, which caused a significant increase instead of a decrease (VAPB p-value=0.0166, VCP p-value=0.0503, and Stathmin-2 p-value=0.0344). VAPB, VCP, and Stathmin-2 all followed a similar trend of having increased (or not significantly changed) concentrations on EVs with the addition of macropinocytosis inhibitors Wortmannin and EIPA (VAPB Wortmannin p-value=0.0236, VCP and Stathmin-2 were non-significant), then having decreases on EVs (or not significantly changed) with the addition of caveolin and clathrin endocytosis inhibitors Genistein (VAPB p-value=0.004), MitMab (VAPB p-value=0.024), and Chlorpromazine (VAPB p-value=0.025), as well as exocytosis inhibitors Manumycin A (Stathmin-2 p-value=0.003) and GW4869 (Stathmin-2 p-value=0.0007) (**Figure 5**). The significant increase with inhibition of the macropinocytosis pathway could mean that an alternative endocytosis pathway is compensating for vesicular turnaround and leading to an increase in VAPB, VCP, and Stathmin-2 when trimer is stabilized. Either ESCRT-dependent or independent exocytosis pathways could be contributing to the trimer altered release as could the caveolin and clathrin endocytosis pathways.

**Figure 4.**
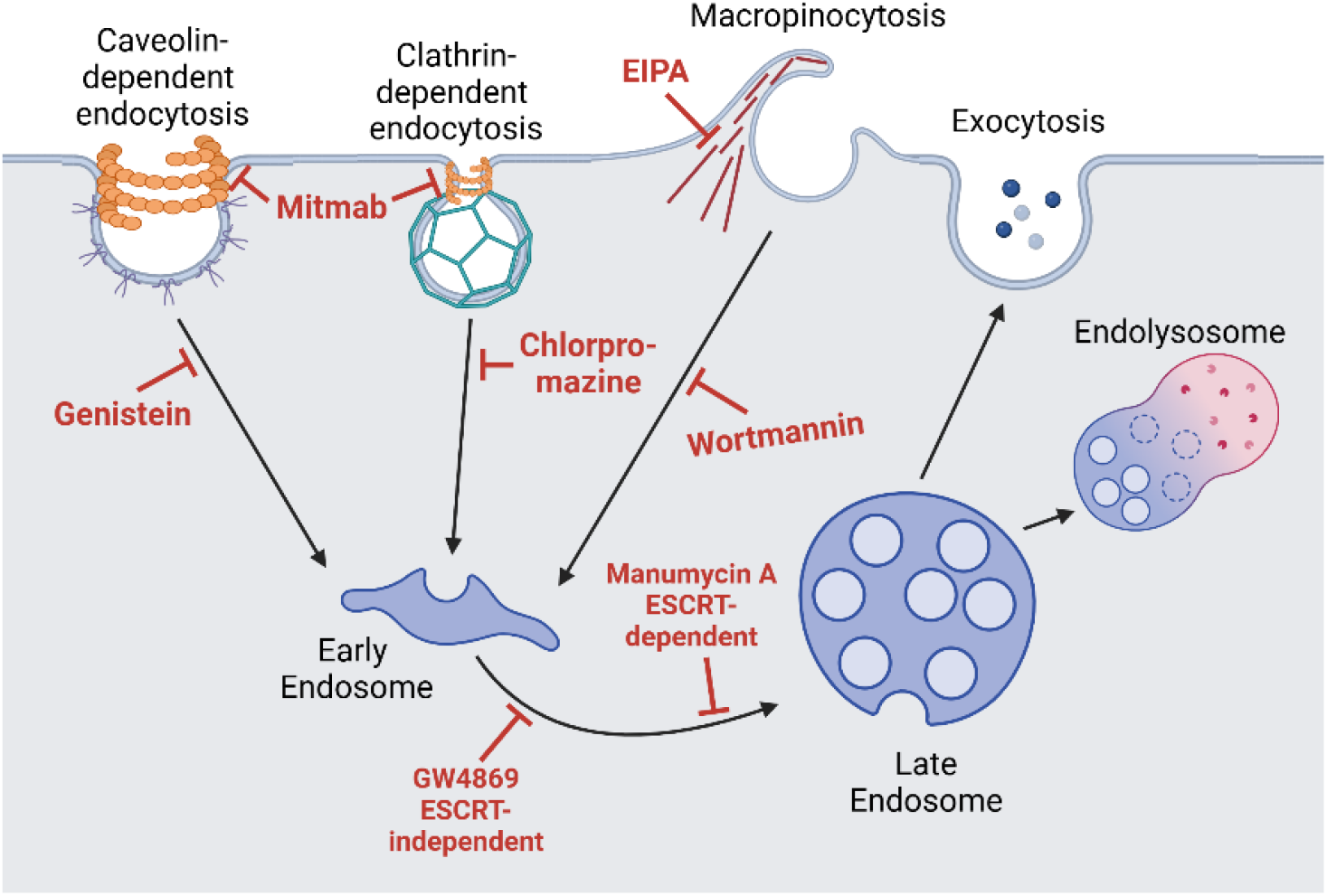
Six different endocytosis and exocytosis inhibitors were utilized to evaluate which EV pathways are being affected by trimeric SOD1 stabilization.

**Figure 5.**
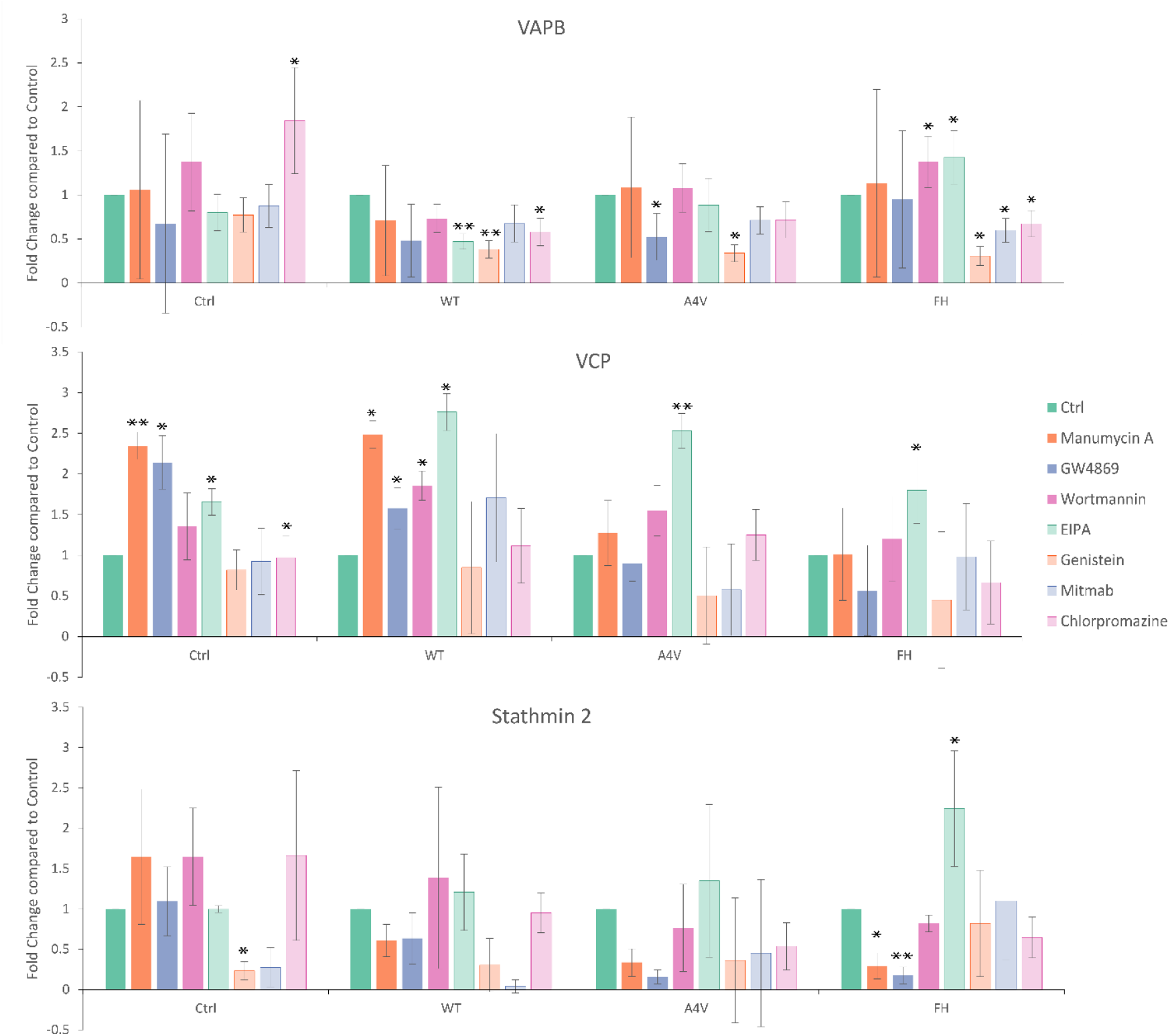
Exocytosis and endocytosis have different effects on VAPB, VCP, and Stathmin-2 levels on CD9+ EVs with trimer stabilization. EIPA increases the level of all three proteins on EVs with trimer stabilization and Genistein lowers the levels. * = p-value <0.05, **= p-value < 0.001.

We also evaluated any changes to twenty-two EV-related proteins on EVs with trimer stabilization. These proteins can be partitioned into six main groups; ER and Golgi function^61^ (Calnexin^62^, Endoplasmic Reticulum protein 72 kDa (ERP72)^63^, Receptor binding cancer antigen expressed on SiSo cells (RCAS1)^64^, Syntaxin 6^65^, and Neural precursor cell expressed, developmentally down-regulated 8 (NEDD8)^66^), lysosomal trafficking (Vesicle-associated membrane protein 7 (VAMP7)^67^ and Lysosome-associated membrane protein 2 (LAMP2)^68,69^), vesicle tethering (Syntaxin-1B, Rab-3 interacting molecule 2 (RIM2), and Syntaxin binding protein 1^70,71^), Vesicle transport/ cytoskeleton (Coronin-1a^72^, Catenin-alpha^73^, Capping protein^74^, CLIP-associated protein (CLASP)^75^, and Septin-7^76^), proteins with direct involvement in endocytosis/ exocytosis (Alix^77^, Caveolin-1, and Cavin-1^78,79^), and Rab family proteins (Rab4, Rab7, Rab9, and Rab11)^40^. Together, these proteins coordinate the formation, cargo loading, and secretion of Evs (**Figure 6A**). RIM2 (p-value= 0.0006 for both FH and A4V) was the only protein testing with a significant decrease on EVs with trimer stabilization. Caveolin-1 (p-value= 0.001) and Cavin-1 (p-value=0.026) had significant increases (**Figure 6B**) on EVs with trimer stabilization. No other proteins tested had significant changes (**Supplementary Figure S4**). Additionally, Cavin-1 oligomers decrease in cell lysate with trimer stabilization, and oligomers of Caveolin-1 increase with trimer stabilization while the overall level of Caveolin-1 was not affected (**Figure 7**). These Caveolin-1 oligomers are obvious in native western blots, but break down rapidly with the addition of 0.1 M Dithiothreitol (DTT) and heat (**Supplementary Figure S5**). Caveolin-1 and Cavin-1 are both inhibited by Genistein and directly interact to regulate Caveolin-dependent endocytosis. Caveolin-1 specifically has been recently connected to ALS^80^; overexpression of Caveolin-1 in an ALS SOD1 mouse model improved neuromuscular function^81^, suggesting a loss of function of Caveolin-1 may be contributing to ALS disease pathology^82^. Ritz et al. previously showed that VCP directly affects ubiquitylation of Caveolin-1 leading to altered Caveolin-1 oligomers similar to what we observed with trimer stabilization^83^. RIM2 helps regulate Rab3, which directly controls vesicle fusion and exocytosis. The vesicle fusion and exocytosis is inhibited by both Manumycin A and GW4869^84^, as well as vesicle transport on the cytoskeleton, which is connected to Stathmin-2^57^. Although there are no direct roles yet known for RIM2 in ALS, RIM2 is crucial to multiple stages of EV release. The combined evidence for the role of the Caveolin-1 pathway, as well as the RIM2-mediated exocytosis pathway in the release of altered EVs with trimeric SOD1 stabilization, suggests multiple pathways are engaged by contributing to a joint undiscovered novel pathway for EV release and spreading in ALS.

**Figure 6.**
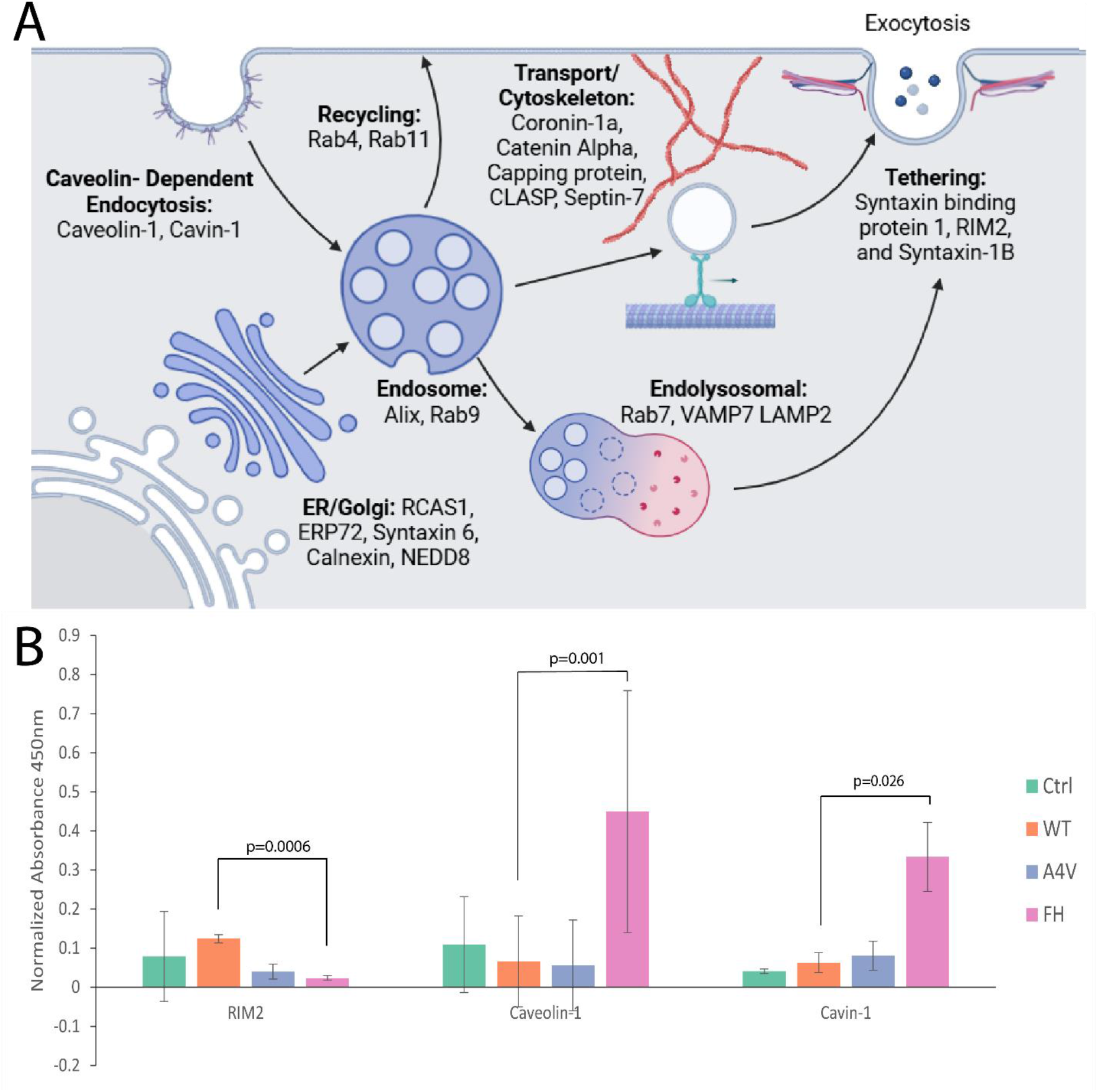
EV-related proteins are affected by SOD1 trimer stabilization. A) Twenty-two different EV-related proteins were evaluated to determine if the levels changed on EVs or in cell lysate with trimer stabilization. B) Only three proteins had significant changes with trimer stabilization; RIM2 decreased (p-value = 0.0006), Caveolin-1 (p-value=0.001) and Cavin-1 (p-value= 0.026) increased.

**Figure 7.**
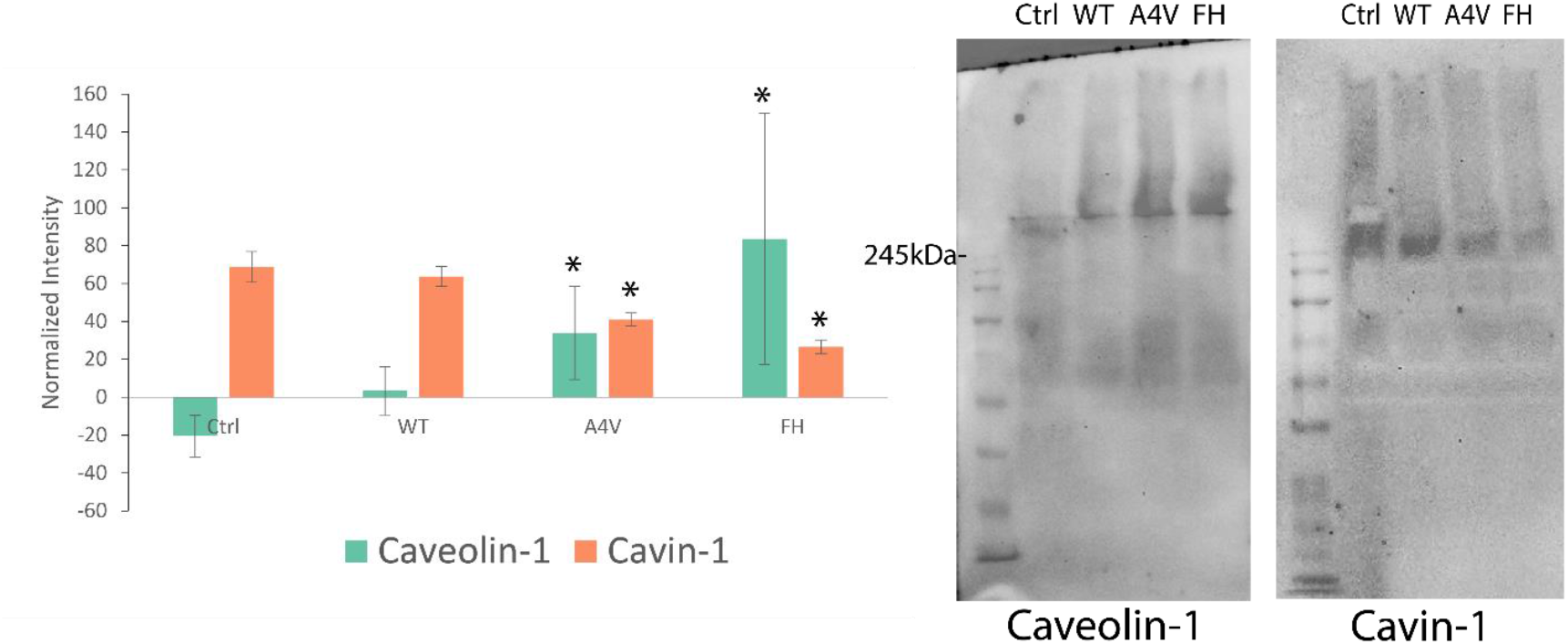
Native western blotting for Caveolin-1 and Cavin-1 display alternate effects in cell lysate. Caveolin-1 oligomers increase with trimer stabilization (WT to A4V p-value= 0.039, WT to FH p-value =0.029), while Cavin-1 oligomers decrease with trimer stabilization (WT to A4V p-value= 0.036, WT to FH p-value =0.015).

## Methods

### Lysate and EV Isolation

We cultured neuroblastoma spinal cord hybrid cells (NSC-34) in six-well plates to 60-80% confluency using 50:50 DMEM:F12 medium supplemented with 10% (v/v) fetal bovine serum and 1% penicillin/ streptomycin. We differentiated the NSC-34 cells into a neuronal phenotype by adding retinoic acid to a final concentration of 10μM^23^ for 48 hours. Once the cells were differentiated, we transiently transfected WT SOD1 or mutants (A4V or FH^23^) using lipofectamine (JetOptimus) in triplicate wells for each mutant (and triplicate control wells that were treated with the transfection reagent and no DNA). Twenty-four hours after transfecting, we switched the medium and allowed the cells to grow for an additional 48 hours, at which point we collected the enriched medium and lysed the cells using RIPA buffer with protease inhibitors. We performed western blotting using the lysed cell samples to confirm transfection of the different ALS protein mutants and to evaluate any changes in other protein levels within the cells due to SOD1 overexpression. Immediately after collecting, we centrifuged the enriched medium at 1,000 rpm to pellet any floating cells. We then transferred the supernatant to an ultracentrifuge tube and centrifuged for two hours at 100,000 x g to pellet the EVs. We resuspended the EV pellet in 1X PBS (50 μL PBS per well of medium) and used it immediately for either ELISA or western blotting. We added RIPA buffer at a 1:1 ratio to any EV pellet samples before western blotting. All samples were normalized before western blotting by calculating the total protein concentration using a Bicinchoninic acid assay (BCA). All western blots were performed using 12% protein gels cast using the Mini-PROTEAN Tetra Handcast System (BioRad) and performed using a Mini-PROTEAN Tetra Vertical Electrophoresis Cell (BioRad) in Tris-glycine running buffer (with or without SDS depending on native or reduced). Reduced samples were prepared using SDS-PAGE loading dye with 0.1 M DTT. We performed all blotting onto PVDF membranes using the BioRad Transblot turbo system before being blocked in 5% BSA solution (diluted in 1X TBST). All blots were incubated in primary antibody overnight at 4 °C and in secondary antibody for 1 hour at room temperature before development using SuperSignal West Pico PLUS Chemiluminescent substrate and imaging using a Chemidoc imager (BioRad). The western blot images were quantified using ImageJ and unpaired t-tests were performed using Python.

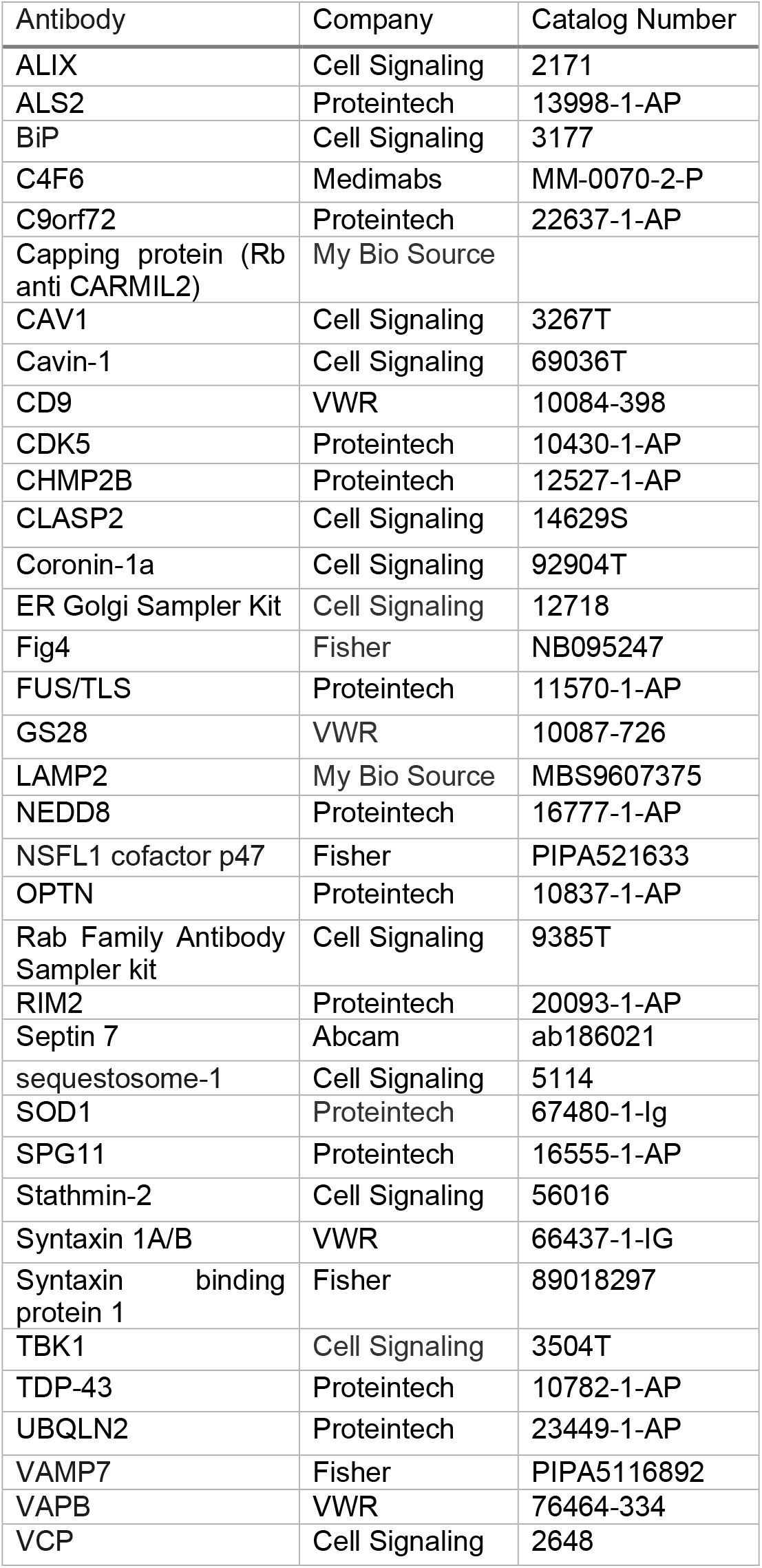
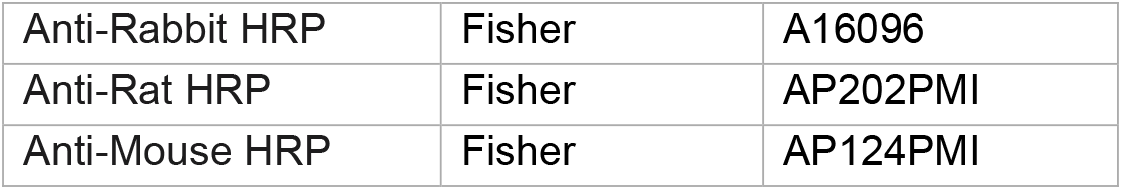

### ELISA

We performed sandwich ELISAs using anti-CD9 as the capture antibody to isolate EVs, then anti-C4F6 (detects trimeric SOD1^30^) or antibodies against other ALS-related proteins FUS, TDP-43, C9orf72, VAPB, SQSTM1, UBQLN2, VCP, OPTN, TBK1, ALS2, CHMP2B, SPG11, Stathmin-2, NEK1, FIG4, NSFL1, and Cdk5 (or other EV-related proteins) as the detection antibody. Rat anti-CD9 was coated at 1 mg/mL and 50 μL/well (diluted in 1X PBS) overnight at 4 C. Following coating we washed the plates 3 times with 1X PBS, then blocked the plate using 1% BSA overnight at 4 °C. The following day, we added isolated EVs (50 μL/ well) to the wells and incubated them for 2.5 hours at 37 °C. Then we washed the plates 3 times with 1X PBS and added primary antibodies (1:1000 for all except C4F6 which was used at 1:250) before leaving the plates at 4 °C overnight. Next, we washed the plates 3 times with 1X PBS and added 50 μL of secondary antibody (anti-Rabbit HRP or anti-Mouse HRP) to each well at 1:1000 for 1 hour at room temperature. After a final wash, we visualized the HRP signal by adding 100 μL TMB substrate (per well) for 5 minutes then stopped the reaction by adding 50 μL of 2N Sulfuric acid. We then quantified the absorbance at 450nm for each well using a SpectraMax i3 plate reader. To account for different concentrations of EVs, we performed an indirect ELISA alongside the sandwich ELISA to quantify the amount of CD9+ vesicles in each sample. We coated ELISA plates with isolated EV samples (diluted 1:5 in 1X PBS) overnight at 4°C. The following morning we washed the plates 3 times with 1X PBS then added Rat anti-CD9 antibody (1:1000) for 1 hour at room temperature. We then washed the plates again and added an anti-Rat HRP secondary antibody for 1 hour at room temperature. After a final washing step, we used TMB substrate to quantify the HRP signal as described previously. We analyzed the ELISA signal by first subtracting the negative control (same process for sandwich ELISA but no EVs added) signal from the sandwich ELISA signal then dividing by the CD9 signal to normalize each well to the total concentration of EV positive EVs. The control samples are from NSC-34 cells transfected with an empty vector. All ELISA sample intensities were normalized to both the signal intensity of triplicate negative control wells (all antibodies added with no sample) as well as to the total concentration of CD9 protein within each sample to normalize for total EV concentration, thus the sample intensity is negative in some cases. The overexpression of SOD1 between the different mutants was confirmed to be consistent using western blotting for total human SOD1 (**Supplementary figure S1**) and none of the significantly changed proteins had any changes in expression within cells with trimer stabilization (**Supplementary figure S2**), the only changes were on EVs. We calculated the p-values for the difference between WT and FH samples as well as the difference between WT and A4V samples using an unpaired two-sample t-test through Python.

### EV mechanism Inhibitors

We grew NSC-34 cells and transfected them with SOD1 mutant constructs following the methods described above. Post-transfection we changed the medium and add DMSO (2 μL/well, vehicle control), Manumycin A (1 μM, ESCRT-dependent exocytosis inhibitor, VWR 102513-528), GW4869 (35 μM, ESCRT-independent exocytosis inhibitor, VWR 103548-206), Wortmannin (25 nM, macropinocytosis inhibitor, VWR 102515-748), EIPA (50 μM, Na/H channel inhibitor, also a factor in macropinocytosis, Sigma A3085-25mg), Chlorpromazine (25 μM, Clathrin-dependent endocytosis inhibitor, VWR 80055-146), Genistein (75 μM, Caveolin-dependent endocytosis inhibitor, VWR 89148-900), or Mitmab (10 μM, Dynamin Inhibitor, Fisher 32-441-1500MG) for 48 hours, then we isolated EVs and performed an ELISA as described above. We determined all inhibitor concentrations by testing five different concentrations within the recommended range of each inhibitor and choosing the highest concentration that did not cause cell death (determined by visualizing the cells on a Keyence BZ-X810 inverted microscope). Alongside the imaging we also performed a lysed EV western blot for the Manumycin A and GW4869 concentration curves for both CD9 and CD63, we chose concentrations that lowered the level of CD63 but did not affect CD9. The fold change was calculated for each inhibitor (compared to the vehicle control) and the error bars were determined by calculating the absolute error using Microsoft Excel.

## Discussion

ALS is currently diagnosed by ruling out other conditions^85^, which makes the process long and challenging. This delay has severe implications for patient outcomes. Without a definitive test, patients often spend months or even years undergoing evaluations before receiving a diagnosis. By that time, the disease has typically progressed, limiting treatment options and reducing the potential benefit of therapeutic interventions. Delated diagnosis contributes to the reduced life expectancy seen in ALS, as patients are often diagnosed too late for meaningful disease-modifying treatments to take effect. Identifying shared factors involved in ALS development could improve our understanding of the disease, guide therapy development, and lead to better diagnostic tools. Proteins found on extracellular components like EVs are promising as clinical biomarkers because they can be measured in biofluids rather than requiring invasive tissue samples^86^. Finding a common pathway responsible for the release of altered EVs in ALS could also reveal new targets to slow or stop disease progression. Many studies have shown that EVs play a role in how ALS spreads in the body^8,31^, suggesting that they are altered in a way that allows the disease to move between cells. Previous research has mostly focused on how common ALS-related mutations affect the spread of misfolded SOD1 on EVs^33^. Other studies have shown that proteins like FUS and TDP-43—both commonly mutated in ALS—can promote the appearance of misfolded SOD1 on EVs^26^. This work takes a new approach by showing that misfolded (trimeric) SOD1 on EVs can, in turn, alter the levels of several ALS-associated proteins on EVs, including Caveolin-1. These findings suggest a network of ALS proteins linked through EV-related disease spread, which were previously thought to act independently.

The field of EV research (especially small EVs like exosomes) has only gained traction within the last decade and there is still very little known about EV formation and release^87^, especially in connection to disease states. Studying EVs can be difficult as methods are still evolving, despite using the same conditions on multiple repeats of the assays included in this work there was still a large amount of variability potentially due to; differences in the passage number of each cell group used, time between ultracentrifugation and isolating the pellet (vesicle pellets typically stayed within the tube when removing the media but would occasionally begin dispersing back into the solution with any slight movement), and vesicle concentrations for the ELISA assays (although we scaled the cell plating to the amount of antibodies being tested for each ELISA, scaling experiments can cause unforeseen changes in the pelleting of vesicles). We accounted for these variables by normalizing all ELISAs to the total concentration of CD9+ vesicles within each sample. We chose CD9 to isolate vesicles since numerous previous studies have shown the colocalization of CD9 with misfolded SOD1 on EVs^35,36^, using a sandwich ELISA approach with a CD9 capture prevents us from losing large amounts of sample while attempting to purify vesicles.

Stabilization of trimeric SOD1 altered the levels of VAPB, VCP, Stathmin-2, SPG11, RIM2, Caveolin-1, and Cavin-1 on EVs—proteins that are all interconnected at various stages of EV biogenesis and release^45,78,84^. Although the mechanistic details of extracellular vesicle (EV) biogenesis remain incomplete, our data suggest these proteins may converge within a common vesicle trafficking and proteostasis pathway that is dysregulated in ALS (**Figure 8**). Supporting this model, we found that VAPB, VCP, and Stathmin-2 levels significantly increased on EVs in the presence of trimeric SOD1, while SPG11 levels decreased, indicating a selective reprogramming of vesicle content. The upregulation of VAPB and VCP—both essential regulators of ER and endolysosomal trafficking^83,88^—suggests that trimeric SOD1 directs these proteins into EV-associated pathways, enhancing the release of proteostasis-related cargo. Macropinocytosis inhibitors (wortmannin and EIPA) further amplified EV levels of VAPB, VCP, and Stathmin-2, implying that when compensatory uptake mechanisms are blocked, SOD1-driven EV secretion is intensified, possibly as a cellular stress response. In contrast, genistein, a Caveolin-1 inhibitor, suppressed all three proteins on EVs, implicating Caveolin-dependent trafficking in this process. Interestingly, Manumycin A and GW4869, which inhibit ESCRT-independent EV release, did not affect VAPB or VCP levels but selectively reduced Stathmin-2, suggesting that distinct EV biogenesis routes may be exploited differently by trimeric SOD1.

**Figure 8.**
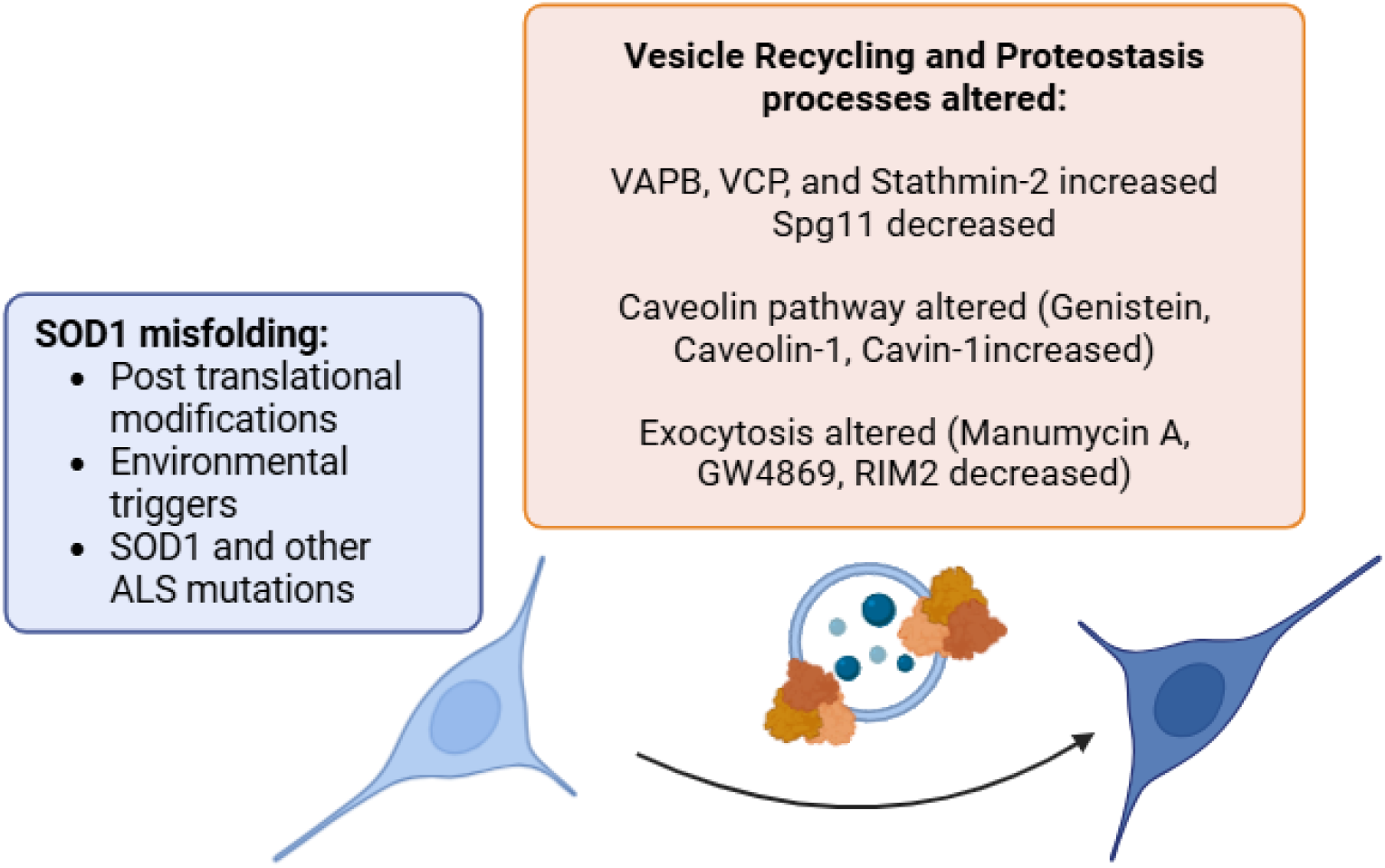
Stabilization of trimeric SOD1 alters EVs by potentially affecting vesicle recycling and proteostasis.

Caveolin-1 and Cavin-1 levels increased on EVs with trimer stabilization, while RIM2, a regulator of synaptic vesicle exocytosis^84^, decreased, indicating a shift in vesicle origin or release mechanism. Within cells, we observed a marked increase in oligomeric Caveolin-1 and a corresponding decrease in Cavin-1 oligomers, consistent with the formation of abnormal Caveolin-rich membrane domains and displacement of Cavin-1. This imbalance may result in membrane deformation, defective vesicle sorting, and altered fusion dynamics. Together, these findings suggest that trimeric SOD1 not only disrupts intracellular proteostasis but also hijacks membrane trafficking machinery, altering EV composition and promoting the spread of pathology. The accumulation of trimeric SOD1 aligns with elevated oligomeric Caveolin-1 within cells, supporting a model in which misfolded SOD1 reshapes membrane domains and reroutes vesicle export pathways. Given the known roles of VAPB, VCP, Caveolin-1, and Cavin-1 in virus budding and vesicle system hijacking^89–92^, a toxic gain-of-function mechanism is likely at play. This viral-like exploitation of the endocytic and exocytic machinery may explain the broad convergence of disrupted pathways in ALS. Our results support the existence of a unified trafficking axis, driven by toxic protein assemblies, that facilitates disease spread and offers novel targets for therapeutic and biomarker development (**Figure 8**).

## Acknowledgment

We acknowledge support from the National Institutes for Health (R35 GM134864), the National Science Foundation (2210963), and the Passan Foundation. Figures were created in https://BioRender.com.

## Author Contributions

B.H. and N.V.D. were the originators and leaders of the project. B.H. carried out all assays. B.H. and N.V.D. wrote the manuscript.

