## Supplemental File for "Novel extracellular vesicle release pathway facilitated by toxic superoxide dismutase 1 oligomers"

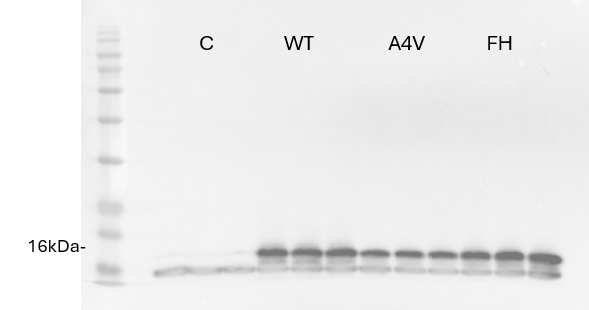

**Figure S1.** Western blots of NSC-34 lysate overexpressing SOD1 WT or mutants (A4V or FH).

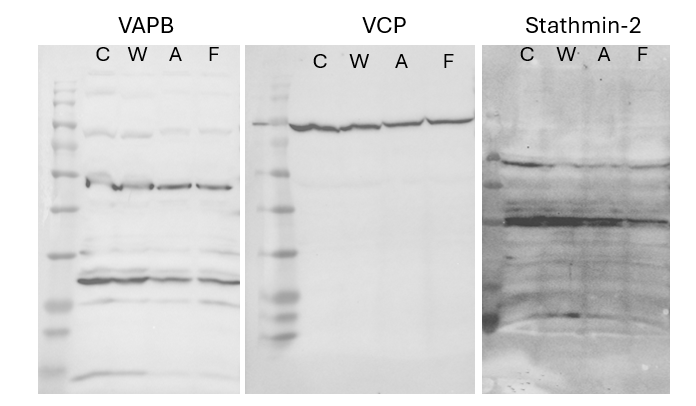

**Figure S2.** VAPB, VCP, Stathmin-2 levels do not significantly change in NSC-34 lysate with SOD1 overexpression (C=Control, W= Wild type, A= A4V, F= FH).

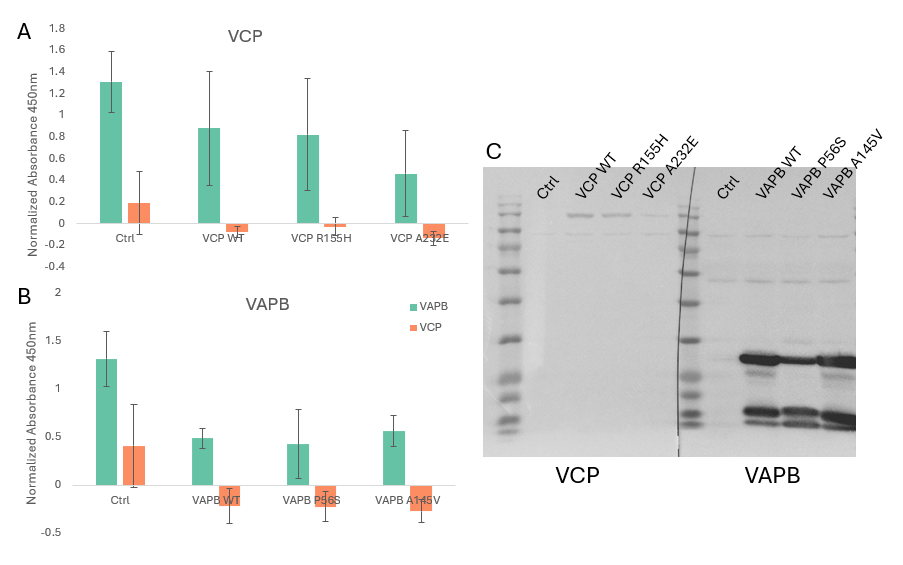

**Figure S3.** Overexpression of VAPB, VCP mutants does not induce the same increases in VAPB and VCP on CD9+ vesicles that is observed with trimeric SOD1 stabilization. Overexpression of the VCP and VAPB mutants in NSC-34 cells was confirmed using western blotting.

Normalized Absorbance 450nm

|  | Ctrl | WT | A4V | FH |
| --- | --- | --- | --- | --- |
| VAMP7 | 0.167692 | 0.0933221 | 0.0152247 | 0.21367 |
| NEDD8 | 0.106141 | 0.08817 | 0.0860994 | 0.0478581 |
| Lamp2 | 0.0660602 | -0.0342157 | -0.00926659 | 0.0244594 |
| Capping | 0.22442 | 0.203585 | 0.109928 | 0.201392 |
| Coronin-1a | 0.189522 | 0.0731456 | 0.0954896 | 0.226037 |
| Septin 7 | -0.0520193 | -0.185765 | -0.122651 | -0.0662381 |
| Syntaxin Binding Protein | -0.0209674 | 0.00568852 | 0.0789731 | 0.107587 |
| Syntaxin 1 | 0.13568 | 0.210972 | 0.173457 | 0.172825 |
| Alix | 0.0416236 | 0.0435492 | 0.0754516 | 0.0301441 |

Normalized Absorbance 450nm

|  | Ctrl | WT | A4V | FH |
| --- | --- | --- | --- | --- |
| RCAS1 | 0.0959554 | 0.167579 | 0.188689 | 0.158928 |
| ERP72 | 0.0289757 | -0.0153106 | 0.0346404 | 0.10969 |
| Syntaxin 6 | -0.0659886 | -0.120427 | -0.121105 | -0.0220202 |
| Calnexin | -0.0345862 | -0.135472 | -0.12611 | -0.0117459 |
| GS28 | 0.186108 | 0.172901 | 0.210487 | 0.110496 |
| CLASP | 0.11399 | 0.0618205 | 0.0305021 | 0.0875591 |
| Rab4 | 0.378304 | 0.245341 | 0.233796 | 0.396324 |
| Rab7 | 0.191066 | 0.0681226 | 0.0742172 | 0.189438 |
| Rab9 | 0.143244 | -0.00769118 | -0.0136519 | 0.075247 |
| Rab11 | 0.11807 | -0.0986613 | -0.135396 | 0.0525074 |

**Figure S4**. Alterations on twenty-two EV-associated proteins were tested on EVs with the stabilization of trimeric SOD1, these nineteen proteins had no significant changes.

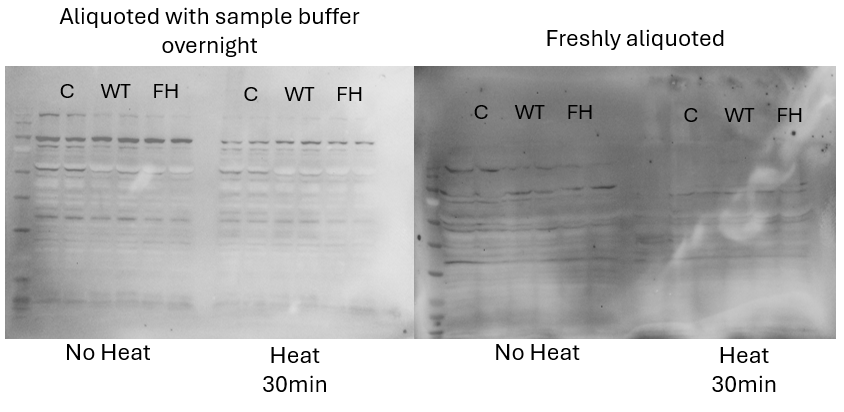

**Figure S5.** Stabilization of trimeric SOD1 (FH) promotes increases in oligomeric Caveolin-1 species that are also observed in native blots. Oligomeric Caveolin-1 breaks down or forms different oligomers after overnight incubation in sample buffer (0.1M DTT) and after heating the samples.
